# Beyond broadband: towards a spectral decomposition of EEG microstates

**DOI:** 10.1101/2020.10.16.342378

**Authors:** Victor Férat, Martin Seeber, Christoph M. Michel, Tomas Ros

## Abstract

Originally applied to alpha oscillations in the 1970s, MS analysis has since been used to decompose mainly broadband EEG signals (e.g. 1-40 Hz). We hypothesized that MS decomposition within separate, narrow frequency bands could provide more fine-grained information for capturing the spatio-temporal complexity of multichannel EEG. In this study using a large open-access dataset (n=203), we decomposed EEG recordings into 4 classical frequency bands (delta, theta, alpha, beta) in order to compare their individual MS segmentations using mutual information as well as traditional MS measures (e.g. mean duration, time coverage). Firstly, we confirmed that MS topographies were spatially equivalent across all frequencies, matching the canonical broadband maps (A, B, C, D). Interestingly however, we observed strong informational independence of MS temporal sequences between spectral bands, together with significant divergence in traditional MS measures. For example, relative to broadband, alpha/beta band dynamics displayed greater time coverage of maps A & B, while map D was more prevalent in delta/theta bands. Moreover, by using a frequency-specificMS taxonomy (e.g. θA, αC), we were able to predict the eyes-open vs closed-behavioural state significantly better using alpha-band MS features compared with broadband ones (80% vs 73% accuracy). Overall, our findings demonstrate the value and validity of spectrally-specific MS analyses, which may prove useful for identifying new neural mechanisms in fundamental research and/or for biomarker discovery in clinical populations.

## 1 Introduction

Multi-channel Electroencephalography (EEG) is a long-established tool for exploring the human brain’s spatiotemporal activities. Microstate (MS) analysis (Michel and Koenig 2018), first introduced by Lehmann (Lehmann 1971) in 1971, takes advantage of EEG’s high temporal resolution to segment EEG signals into short successive periods of time characterized by metastable scalp topographies. Initially applied to narrow-band alpha oscillations (8 -12 Hz)(Lehmann 1971), microstate analysis is nowadays usually performed on broadband EEG signals (1 – 40Hz) (Michel and Koenig 2018; Zanesco et al. 2020). Historically, only a limited number of studies (Javed et al. 2019; Merrin et al. 1990; Musaeus et al. 2020) have focused on applying MS analysis to the traditional frequencies associated with cortical oscillations (e.g. delta, theta, alpha, beta etc.). For example, in the 1990’s, Merrin et al (Merrin et al. 1990) were the first to report on a significant difference in MS segments between schizophrenic patients and controls specifically in the theta EEG band. On the other hand, more recent work in healthy subjects found that MS dynamics were independent of EEG power fluctuations across the frequency spectrum (Britz, Van De Ville, and Michel 2010), which technically supported the rationale for performing broadband MS analysis. Neuroimaging studies have nevertheless emerged showing that anatomically-distinct cortical regions display different dominant EEG frequencies, with occipito-parietal regions more active in the alpha band, and prefrontal regions being biased more toward delta or theta power (Groppe et al. 2013; Keitel and Gross 2016; Mellem et al. 2017). Moreover, ongoing cortical dynamics have been reported to fluctuate from a local resting/idling alpha oscillatory state to task-specific active mode(s) dominated by other rhythms (e.g. theta (Ribary, Doesburg, and Ward 2017), gamma (Hipp, Engel, and Siegel 2011)). As a consequence, cortical regions could combine different frequencies for integrating/segregating information across large-scale networks, a phenomenon termed “oscillatory multiplexing” (Akam and Kullmann 2014). Finally, of more clinical significance, a growing body of work has indicated abnormal EEG spectral power in distinct frequencies across cortical regions in a variety of brain disorders (Ros et al. 2014; Schulman et al. 2011). Therefore, given that different spatial topographies uncovered by MS analysis imply anatomically-distinct cortical generators (according to the forward-model of EEG generation (Michel and Koenig 2018)), it is reasonable to hypothesize that distinct MS topographies may display different spatial and/or temporal profiles across the frequency spectrum.

To investigate this question as well as gain a deeper understanding of frequency-specific MS signature(s), we sought to explicitly decompose MS spatiotemporal dynamics within *discrete, narrow-band frequency bands* (i.e. delta, theta, alpha, and beta), with the aim of comparing them to the classical analysis of the broadband signal.

Here, we employed a validated, open-source dataset (Babayan et al. 2019) of resting-state EEG recordings from 203 healthy subjects during both eyes opened and eyes closed conditions. These were then filtered in the classical EEG bands (delta: 0-4 Hz, theta: 4-8 Hz, alpha: 8-12 Hz, beta: 15-30 Hz) to obtain band specific signals. These narrow-band signals, in addition to the broadband (1-30 Hz) signal, were then independently subjected to standard microstate analysis (Pascual-Marqui, Michel, and Lehmann 1995). Map topography, mean duration, occurrence, time coverage, and global explained variance (GEV) were used as quantitative measures of spatiotemporal microstate dynamics. In summary, and using spatial correlation analysis, we firstly demonstrate remarkably similar microstate topographies across frequencies, closely matching the classical broadband maps. Interestingly, however, we observed strong informational independence of microstate sequences between frequencies, in addition to significant differences in established measures of temporal dynamics (mean duration, occurrence, and time coverage).

In conclusion, our results support a more diverse, frequency-specific application of microstate analysis compatible with the narrow-band MS analyses of early pioneers (Lehmann 1971; Merrin et al. 1990). We anticipate this approach to provide a more fine-grained spectral information not visible to the standard broadband analysis, for example in the identification of biomarkers in clinical populations or for understanding the mechanisms underlying EEG microstates.

## 2 Methods

### 2.1 Dataset

EEG recordings were obtained from 203 anonymized participants enrolled in the Mind-Brain-Body study (Babayan et al. 2019). Detailed protocol and inclusion criteria are reported in (Babayan et al. 2019). The overall sample consisted of 227 participants divided into 2 groups: the younger adults group with participant age ranging between 20 and 35 years (N = 153, 45 females, mean age = 25.1 years, SD = 3.1) and an older adults group with age ranging between 59 and 77 years (N = 74, 37 females, mean age = 67.6 years, SD = 4.7). Medical and psychological screening was conducted on all participants at the Day Clinic for Cognitive Neurology of the University Clinic Leipzig and the Max Planck Institute for Human and Cognitive and Brain Sciences in order to include only healthy patients. The study protocol was approved by the ethics committee of the University of Leipzig (reference 154/13-ff). Data were obtained in accordance with the Declaration of Helsinki.

### 2.2 Recordings

Resting state EEGs were recorded using 61 scalp electrodes (ActiCAP, Brain Products GmbH, Gilching, Germany), and one additional VEOG electrode for recording right eye activity. All electrodes were placed according to the international standard 10–20 extended localization system with FCz reference, digitized with a sampling frequency of fs=2500 Hz,an amplitude resolution of 0.1 microV, and bandpass filtered between 0.015Hz and 1 kHz. The ground was located at the sternum and scalp electrode impedance was kept below 5 KΩ. Recordings took place in an electrically shielded and sound-attenuated EEG booth. Here, 60s blocks alternated between eyes open (EO) and eyes closed (EC) conditions for a total recording of 16 min (8 blocks EC, 8 blocks EO, starting with EC). During the EO condition, participants were asked to stay awake while fixating their eyes on a black cross presented on a white background.

### 2.3 Prepocessing

The prepossessing steps are extensively described in (Babayan et al. 2019), which we summarize below. All EEG recordings were down-sampled from 2500 to 250 Hz and filtered between 1 and 45Hz (8th order, Butterworth filter). Blocks sharing the same condition were concatenated leading to the creation of 2 datasets per subject. After visual inspection, outlying channels were rejected and EEG segments presenting noise and/or artefacts were removed (except eye movements and eye blinks that were kept for further prepossessing). PCA was used to reduce data dimensionality, by keeping PCs (N≥30) that explain 95% of the total data variance. Then, independent component analysis (ICA) was performed using the Infomax (runica) algorithm. Components reflecting eye movement, eye blink or heartbeat related artefacts were removed.

Before performing microstate analysis, the following additional prepossessing steps were conducted using MNE-python (Gramfort et al. 2013): missing/bad channels were interpolated using spherical spline interpolation, the reference was re-projected to average and recordings were down-sampled to 100Hz. Finally, each recording was filtered into broadband plus the 5 traditional EEG frequency bands: broadband (1-30 Hz), delta (1-4 Hz), theta (4-8 Hz), alpha (8-12 Hz), beta (15-30 Hz). Filter design consisted of a two-pass forward and reverse, zero-phase, non-causal band-pass FIR filter with the following parameters.

- **Broadband (1-30 Hz)**: - Lower passband edge: 1.00 - Lower transition bandwidth: 1.00 Hz (−12 dB cutoff frequency: 0.50 Hz) - Upper passband edge: 30.00 Hz-Upper transition bandwidth: 7.50 Hz (−12 dB cutoff frequency: 33.75 Hz) Filter length: 331 samples (3.310 sec)
- **Delta (1-4 Hz)**: - Lower passband edge: 1.00 - Lower transition bandwidth: 1.00 Hz (−12 dB cutoff frequency: 0.50 Hz) – Upper passband edge: 4.00 Hz - Upper transition bandwidth: 2.00 Hz (−12 dB cutoff frequency: 5.00 Hz) - Filter length: 331 samples (3.310 sec)
- **Theta (4-8 Hz)**: - Lower passband edge: 4.00 - Lower transition bandwidth: 2.00 Hz (−12 dB cutoff frequency: 3.00 Hz) - Upper passband edge: 8.00 Hz - Upper transition bandwidth: 2.00 Hz (−12 dB cutoff frequency: 9.00 Hz) - Filter length: 165 samples (1.650 sec)
- **Alpha (8-12 Hz)**: - Lower passband edge: 8.00 - Lower transition bandwidth: 2.00 Hz (−12 dB cutoff frequency: 7.00 Hz) - Upper passband edge: 12.00 Hz - Upper transition bandwidth: 3.00 Hz (−12 dB cutoff frequency: 13.50 Hz) - Filter length: 165 samples (1.650 sec)
- **Beta (15-30 Hz)**: - Lower passband edge: 15.00 - Lower transition bandwidth: 3.75 Hz (−12 dB cutoff frequency: 13.12 Hz) - Upper passband edge: 30.00 Hz - Upper transition bandwidth: 7.50 Hz (−12 dB cutoff frequency: 33.75 Hz) - Filter length: 89 samples (0.890 sec)

For all filters, a Hamming window with 0.0194 passband ripple and 53 dB stopband attenuation was used to reduce border effects.

### 2.4 MS segmentation

#### 2.4.1 Segmentation

Microstate segmentation was applied to each combination of frequency band (broadband, delta, theta, alpha, beta) x behavioural condition (EO, EC) leading to the computation of 10 optimal clusters using the methodology described below. First, local maxima of the Global Field Power (GFP) known to represent portions of EEG data with highest signal to noise ratio (Koenig and Brandeis 2016), were extracted from each individual recording. Then, 20 epochs of 500 time points randomly drawn from the previous extraction were submitted to a modified k-means cluster analysis using the free academic software Cartool (Brunet, Murray, and Michel 2011). For each number of cluster centers K ranging from 1 to 12, 50 k-means initialisations were applied to each epoch. The initialisation with highest global explained variance (GEV) was selected and kept for further processing. A meta-criterion (Bréchet et al. 2019) was used to choose the optimal number of cluster centers k for each epoch. Individual optimal clusters were then merged within conditions and within frequencies to form 10 groups of 4060 clusters. Each group was then randomly re-sampled into 100 epochs of 5000 time points, and submitted to the same clustering algorithm (50 initialisations, with meta criterion selection), leading to the extraction of 100 optimal clusters per group. Finally, these 100 clusters were submitted to the modified K means clustering algorithm to extract, for each number of cluster centroids k, a set of maps which best represent the spatiotemporal variance of frequency specific EEG data within each condition.

#### 2.4.2 Selection of “common” MS maps

Given that we found high spatial correlations between MS maps across all frequencies and EO/EC conditions, we fitted the broadband maps directly to all the frequency bands in order to have a common reference. This may be considered a heuristic approach for the sake of simplicity. An alternative approach we explored was to perform subject-level (i.e. 1st level) clustering on all data concatenated *within-subject* (across frequencies, and/or conditions), followed by group-level (i.e. 2nd-level) clustering. We found this to once again produce identical maps to the broadband decomposition. This method could theoretically be used to find the most “common” clusters across different datasets, in case of variable k-means outputs (e.g. visually similar MS maps at different k-values). Since it is beyond the scope of this paper, we leave it to future studies to validate this method more rigorously.

#### 2.4.3 Back-fitting of MS Maps

The common topographic maps selected above were then assigned to every time point from all individual recordings using the traditional MS back-fitting method (Van De Ville, Britz, and Michel 2010). First, the spatial correlation was computed between every timepoint and map. Using the so called ‘winner takes all’ algorithm, each timepoint was labelled according to the map with which it shared the highest absolute spatial correlation. Timepoints were labelled as “non-assigned” when the absolute spatial correlation was below r < 0.5 threshold. To ensure temporal continuity of MS segmentation, a smoothing step (Brunet et al. 2011; Pascual-Marqui et al. 1995) was applied. Finally, segments with duration shorter than 3 samples (30ms) were assigned to neighbouring segments using the following rule: the segment was split into two parts, where each part was assigned to the neighbouring segment with the higher spatial correlation. With backfitting completed, we extracted 4 spatiotemporal parameters for each microstate map, namely:

- **Global explained variance** (Gev) described as the sum of variances of the original recording explained by the considered microstate map weighted by the Global Field Power at each moment in time. Units are percentages (%) between 0 and 1.
- **Mean duration** (MeanDurs), defined as the mean temporal duration of segments assigned to each MS map. Units are in seconds (s).
- **Time coverage** (TimeCov) is the ratio of time frames assigned to each MS map relative to the total number of time frames from the recording. Results are Units are percentages (%) between 0 and 1.

### 2.5 Adjusted Mutual information score

Scikit-learn (Pedregosa et al. 2011) implementation of the adjusted mutual information score (AMI) (Vinh, Epps, and Bailey 2010) was used to quantify the mutual information (MI) shared between different MS temporal segmentations, whilst simultaneously accounting for random overlap due to chance. This metric, bounded between 0 and 1, is used to evaluate the statistical (in)dependence of two variables. In our case, AMI is estimated between the symbolic sequences of two different microstate segmentations (e.g. ABDCADB vs ABDBDAC). A high score (approaching 1) indicates that the two segmentations agree on the temporal order of all labels while a low score (approaching 0) indicates that the segmentations’ labels are not temporally aligned. We selected the corrected version of this metric in order to control for the impact of differences in label distribution due to chance (for example differences in overall time coverage between labels).

### 2.6 Statistics

Statistical analyses were performed on the 4 main spatiotemporal parameters (Global explained variance, Mean spatial correlation, Mean duration, Time coverage). Tests were conducted using a two sided permutation test for equality of means on paired samples (same subject, either between condition, either between frequencies) under the H0 hypothesis that both frequency (i.e. condition) share the same mean against the alternative H1 that the distributions come from two different populations. P-values were estimated by simulated random sampling with 10000 replications. As a large number of statistical tests were carried out without specific pre-planned hypotheses (Armstrong 2014), P values were corrected for multiple comparisons using the Bonferroni method. Corrected P-values are reported in the Results section, as well as the observed means (m) of both samples together with observed standard deviations. Effect sizes are reported as the standardised difference of means using Cohen’s d (d).

### 2.7 Neurobehavioral prediction models

#### 2.7.1 Model

Linear Support Vector Classification (SVC) with ‘l2’ norm penalization and squared hinge loss function were used to discriminate EO vs EC states using MS parameters pertaining to broad- as well as narrow-band EEG activity. Models were trained using 15 frequency specific features correspond to the 3 spatiotemporal MS metrics (Gev, MeanDurs, TimeCov) of each maps (A,B,C,D,C’) of a given frequency. Band specific prediction models were fitted with standardized features (after removing their respective mean and scaling them to unit variance).

#### 2.7.2 Evaluation

As suggested by Bouckaer et al. (Bouckaert n.d.), 10 times repeated 10 fold cross validation tests were used to assess classification results. For each of the 100 runs, 9 folds of 20 subject’s features each were used to train the model while 1 fold of 20 subject’s features was used to evaluate the 3 diagnostic metrics:

- **Accuracy**: defined as number of the number of correctly predicted samples out of all the testing set. **The operating characteristic (ROC)** which the plot of true positive rate as a function of the false positive rate. It is used to illustrate the classification trade-off for different discrimination threshold.
- **Area under the curve (AUC)** defined as the area under the ROC curve: it is an aggregate measure of performance for all possible classification thresholds. AUC values are in the range of 0 to 1. A model with 100% error in its predictions has an AUC of 0.0. If all its predictions are correct, its AUC is 1.0.

95% confidence intervals were evaluated on each of the 3 metric distributions with 10 degrees of freedom.

Statistical comparison between models were conducted on both accuracy and AUC using a two-tailed permutation test for equality of means on paired samples under the H0 hypothesis that the alpha-band (8-12 Hz) had a greater mean of the classification metric than the broadband (1-30 Hz). P-values were estimated by simulated random sampling with 10000 replications.

## 3 Results

### 3.1 Spatial Similarity of Microstate Maps

**Figure 1** illustrates the topographic results of MS segmentations in the different conditions and frequency bands. After visual inspection of optimal clusters at different cluster numbers (k), we identified that a value of k=5 revealed five MS topographies that were similar across all EEG bands and behavioural conditions, consistent with recent findings from our laboratory (Bréchet et al. 2019, 2020; D’Croz-Baron et al. 2019). MS maps were designated in line with the canonical prototypes from the literature and their respective symbols, featuring a left-right orientation (A), a right-left orientation (B), an anterior-posterior orientation (C), fronto-central maximum (D) and occipito-central (C’)maximum.

**Figure 1:**
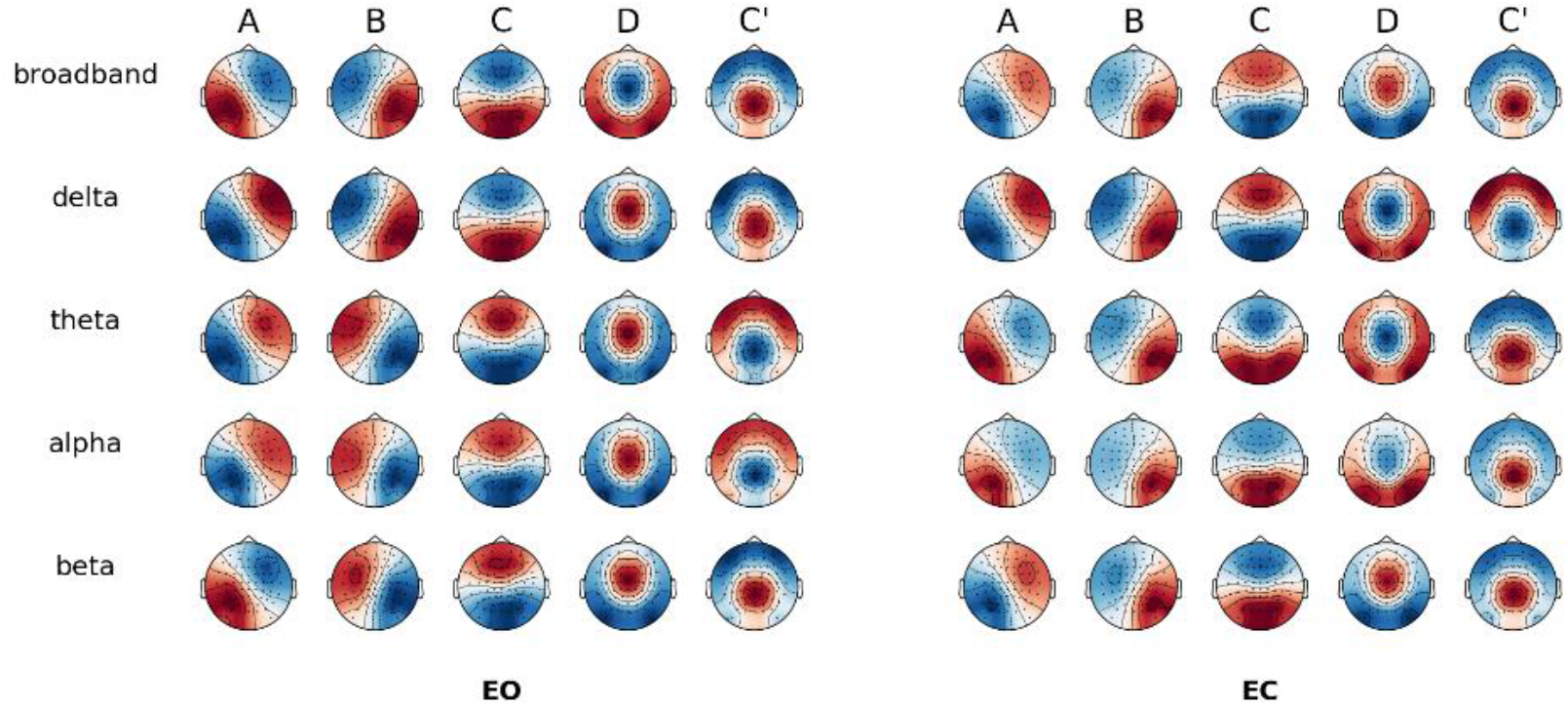
Spatial correlation between MS topographies across behavioural conditions. Global cluster centroids of each frequency band within eyes-open (EO) or eyes-closed (EC) condition. Note that map polarity inversion is ignored in the classical analysis of spontaneous EEG.

Given the additional frequency dimension, we labelled the MS maps firstly according to the Greek letters traditionally used for narrow-band EEG (i.e. δ, θ, α, β) and then the Latin alphabet for the canonical map symbols (i.e. A, B, C, D) . For example, *α*A denoted the left-right diagonal map from the alpha band (*α*) segmentation, and *δ*C the anterior-posterior map from the delta band (*δ*) segmentation. The broadband segmentation was designated with the prefix ‘bb’.

As shown in **Figure 2**, when comparing topographies between broadband and each narrow-band (i.e. the diagonal entries in the correlation matrix), all spatial correlations were *r* > 0.98. Consequently, we fitted the broadband maps directly to all the frequency bands in order to have a common reference.

**Figure 2:**
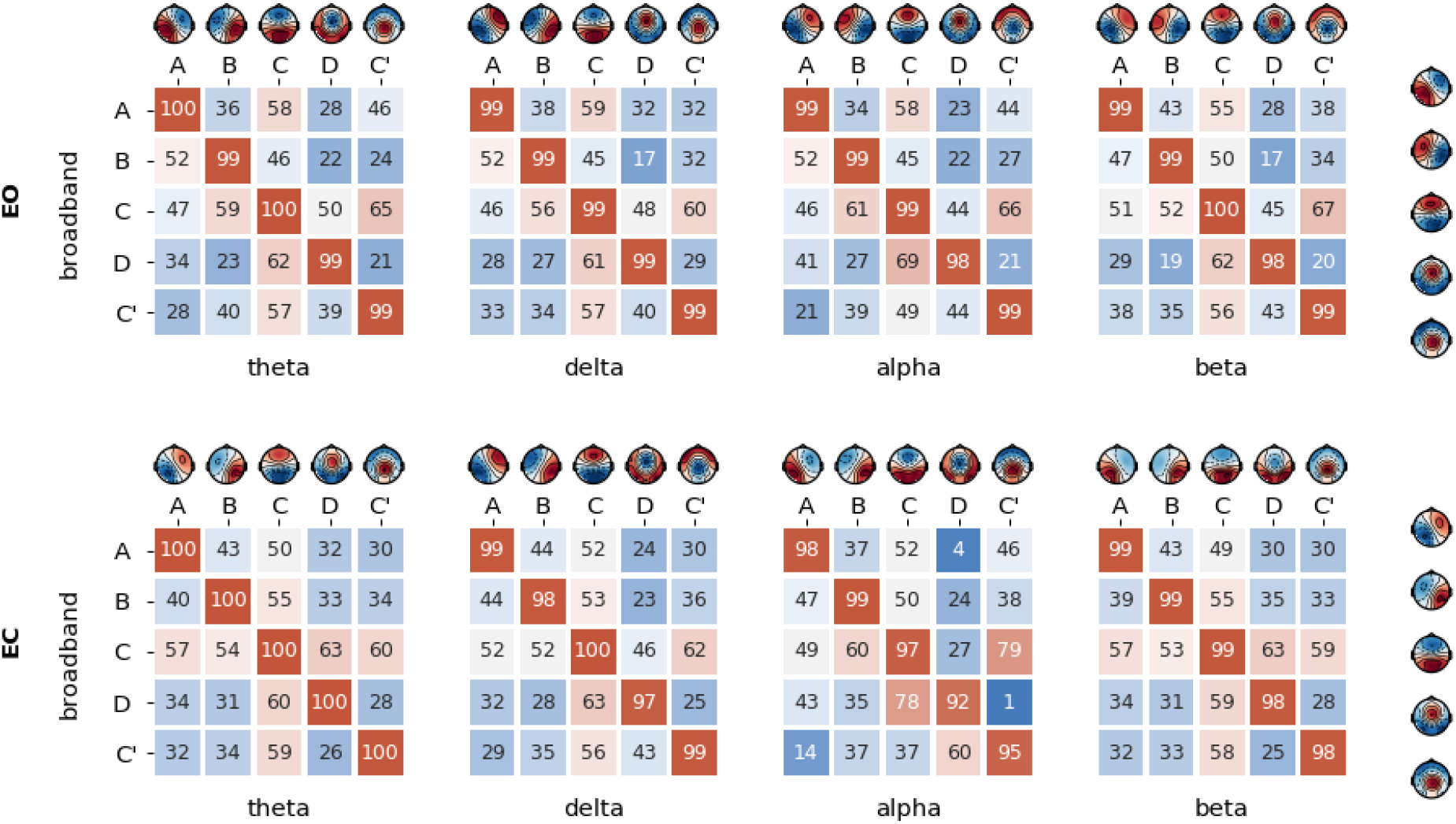
Spatial correlation between MS topographies across frequencies. Spatial correlation of cluster centers of each sub-frequency bands compared to broadband for eyes opened (EO) and eyes closed (EC) condition.

We similarly observed common MS maps when comparing broadband topographies between EO and EC conditions (**Suppl Figure 1**), with all intraclass spatial correlations exceeding *r* > 0.98, thus providing justification for comparing microstate parameters between behavioural conditions while fitting condition specific broadband maps.

### 3.2 Mutual Information of Microstate Sequences

Briefly, Adjusted Mutual Information (AMI, bounded between 0 and 1) is an index of how similar two separate MS segmentations are, by estimating the degree of shared information (i.e. the number of time points assigned with the same MS) between their symbolic sequences (e.g. ABCD vs ABDA). The ‘adjusted’ aspect ensures the measure is unbiased for symbolic overlap(s) due to chance (Vinh et al. 2010). Higher AMI (approaching 1) indicates nearly identical MS temporal sequences, while lower AMI (approaching 0) indicates temporally independent sequences. (I.e. low overlap)

As shown in **Figures 3**, the AMI between broadband and narrow-band segmentations in the EO condition showed a value of s = 0.06 for delta, s = 0.03 for theta, s = 0.05 for alpha, and s = 0.01 for beta. These values are surprisingly low and we can conclude that the broadband segmentation is comparatively independent of the narrow-band one. Similar conclusions of temporal independence can be made by examining the AMI between the narrow-bands *themselves*, with a maximum AMI value between theta and alpha bands (EO: s = 0.006, EC: s = 0.009), and a minimum AMI value of s = 0.001 for non-adjacent EEG bands (delta-alpha, delta-beta, theta-beta)

**Figure 3:**
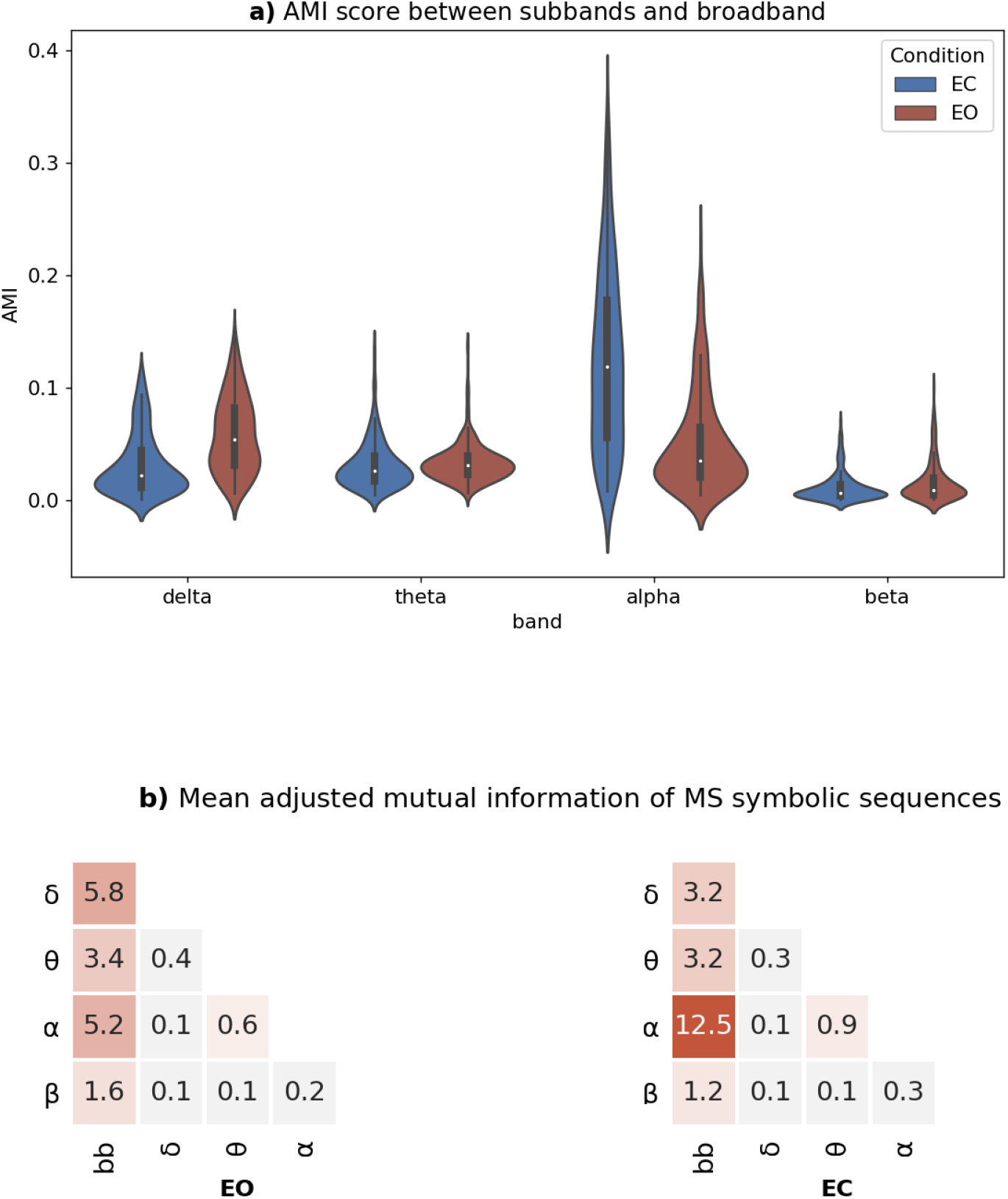
Adjusted mutual information of MS symbolic sequences. **a)**Mean adjusted mutual information (AMI) is depicted, for each behavioural condition (EC: eyes-closed, EO: eyes-open) across n = 203 subjects. **b)**Mean adjusted mutual information of MS symbolic sequences between all broadband and narrow-band combinations. Mean (n = 203 subjects) adjusted mutual information (AMI) for all frequency pairs.

As a sanity check, when inspecting the EO vs EC transition, the shared information with broadband decreased for the delta band (s = 0.03) but increased for the alpha band (s = 0.12). The latter is in line with expectations, as alpha oscillations are known to increase considerably during eye closure, which would amplify their contribution to the broadband signal and consequently their shared dynamics.

### 3.3 Between-frequency comparison of classical MS measures

Here we tested for significant differences between broadband and narrow-band filtered EEG in the classical MS measures: Global explained variance (Gev), Time Coverage (TimeCov), and Mean Duration (MeanDurs). This was done by conducting paired t-tests between broadband (bb) and respective narrow-band (delta to beta) MS measures across all n=203 subjects. For each MS measure and MS map, results were visualised using heat-plots as the narrow-band *absolute difference* from the broadband mean value. Here, a red/blue background indicated significant positive/negative differences at p < 0.05, while a white colour indicated non-significant differences at p > 0.05. Exact p-values and effect sizes are reported in Table 2 of the Supplementary Results.

#### Global Explained Variance (Gev)

**Figure 4** illustrates the percent differences in global explained variance (Gev) between each narrow-band vs broadband, demonstrating the presence of specific “fingerprints” between frequencies (rows) or MS maps (columns)., MS segmentation of **delta** band activity showed significantly higher GEV across most MS maps in both EO (+16 %) and EC (+ 16 %) conditions, compared to the broadband segmentation. A similar but less pronounced effect was found for the theta segmentation, suggesting that low-frequency EEG fluctuations in the 1-8 Hz (delta-theta) range may be accounted for more parsimoniously using the 5 canonical MS maps than 1-30 Hz (broadband) activity.

**Figure 4:**
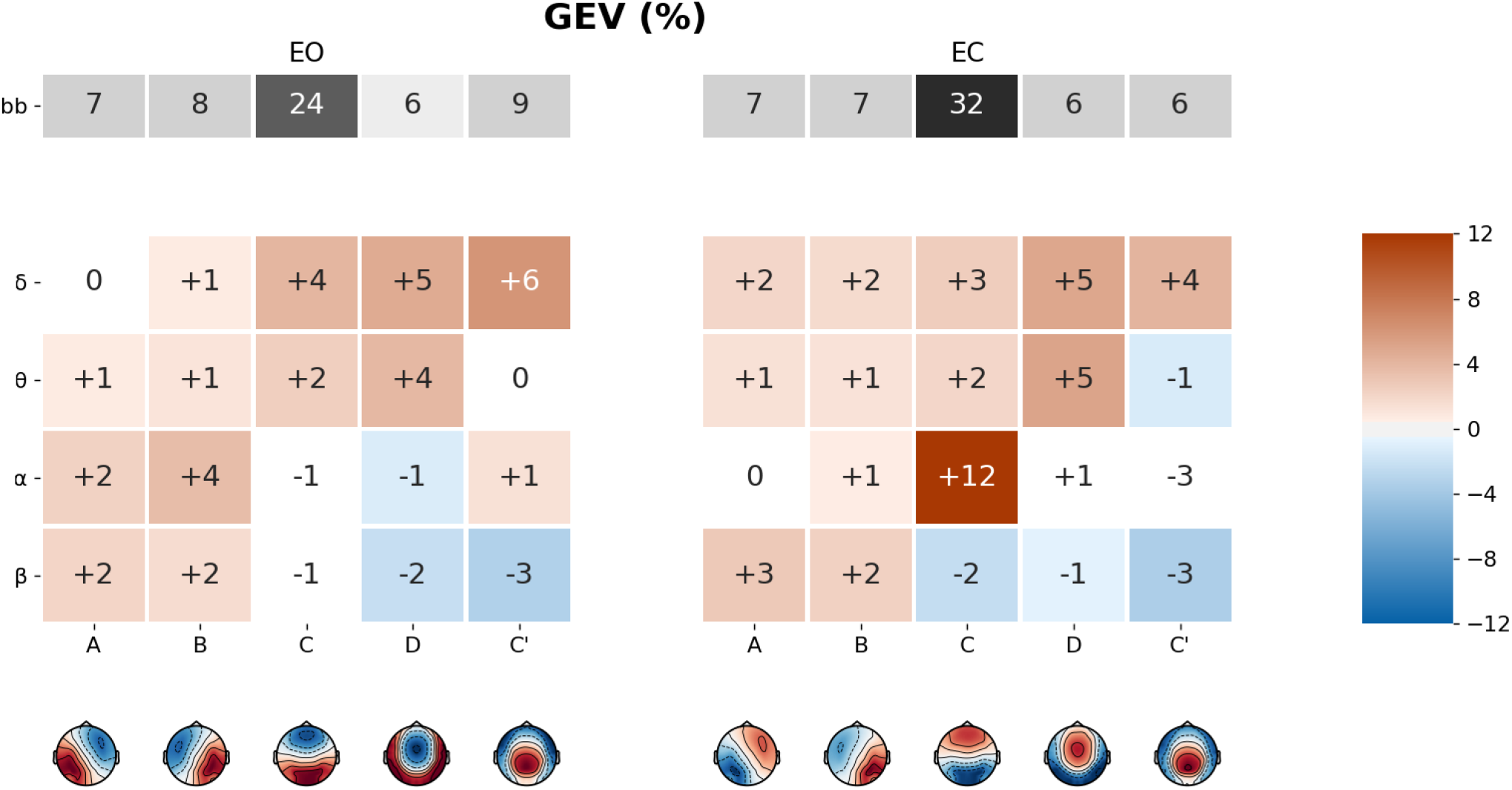
Microstate map differences in Global Explained Variance. (Gev) between broadband and narrow-band filtered EEG, for eyes open (EO) and eyes closed (EC) conditions. The first row (in grey) represents the mean Gev of the broadband segmentation. Coloured rows represent the mean Gev difference between the narrowband (δ, θ, α, β) and broadband segmentations, for each microstate (A – C’). Significant differences have red/blue backgrounds, while non-significant ones have a white/grey background.

The profile of the **alpha** band segmentation depended on behavioural condition. During EC, microstate C had significantly increased Gev (+ 12%) relative to broadband, indicating that distinct MS topographies have stronger behavioural specificity at narrow-band frequencies (see section *Application to Behavioural Classification*). Conversely, during EO, microstates A (+ 2%) and B (+ 4%), expressed significantly more Gev compared to broadband.

Conversely, **or the beta** band, Gev was significantly greater compared to broadband for maps A (EO: + 2%, EC+ 3%) and B (EO: + 2%, EC: + 2%) while it was significantly reduced for maps D (EO: - 2%, EC: - 1%) and C’ (EO: - 3%, EC: - 3%) . Microstate C explained less variance compared to broadband but the comparison was significant only for EC (− 2%).

#### 3.3.1 Time Coverage (TimeCov)

During MS segmentation, each time-point is assigned to only *one* of the five canonical MS topographies through a winner-takes-all process (i.e. the map with the highest spatial correlation with that time-point *wins*). Time Coverage (TimeCov) refers to the *average* prevalence of each MS map over the whole EEG recording **(Figure 5)**, expressed as a percentage (i.e. number of time-points assigned to a particular MS map divided by the total number of time-points).

**Figure 5:**
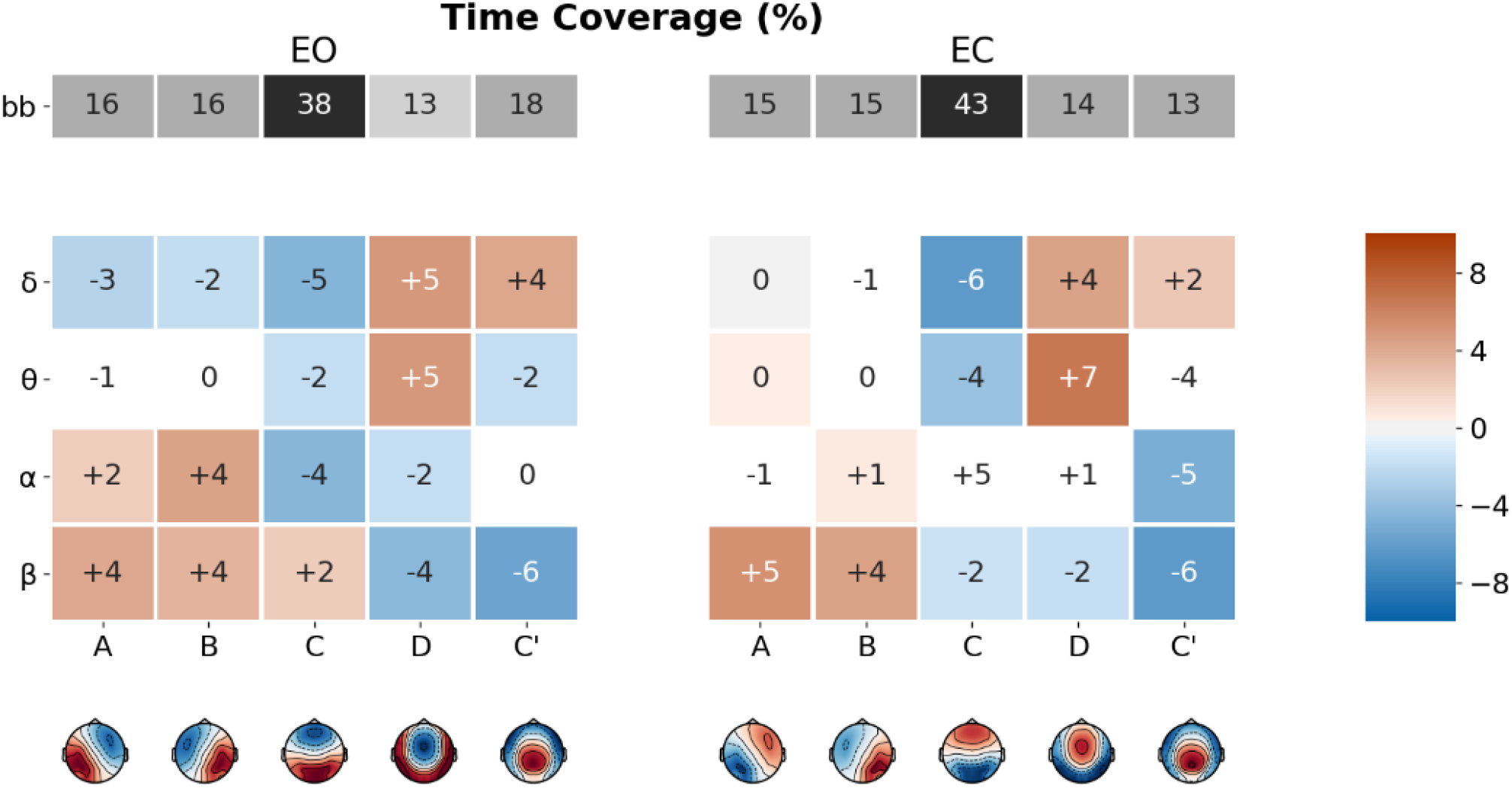
Microstate map differences in Time Coverage. (TimeCov) between broadband and narrow-band filtered EEG, for eyes open (EO) and eyes closed (EC) conditions. The first row (in grey) represents the mean TimeCov of the broadband segmentation. Coloured rows represent the mean TimeCov difference between the narrowband (δ, θ, α, β) and broadband segmentations, for each microstate (A – C’). Significant differences have red/blue backgrounds, while non-significant ones have a white/grey background.

**Figure 6:**
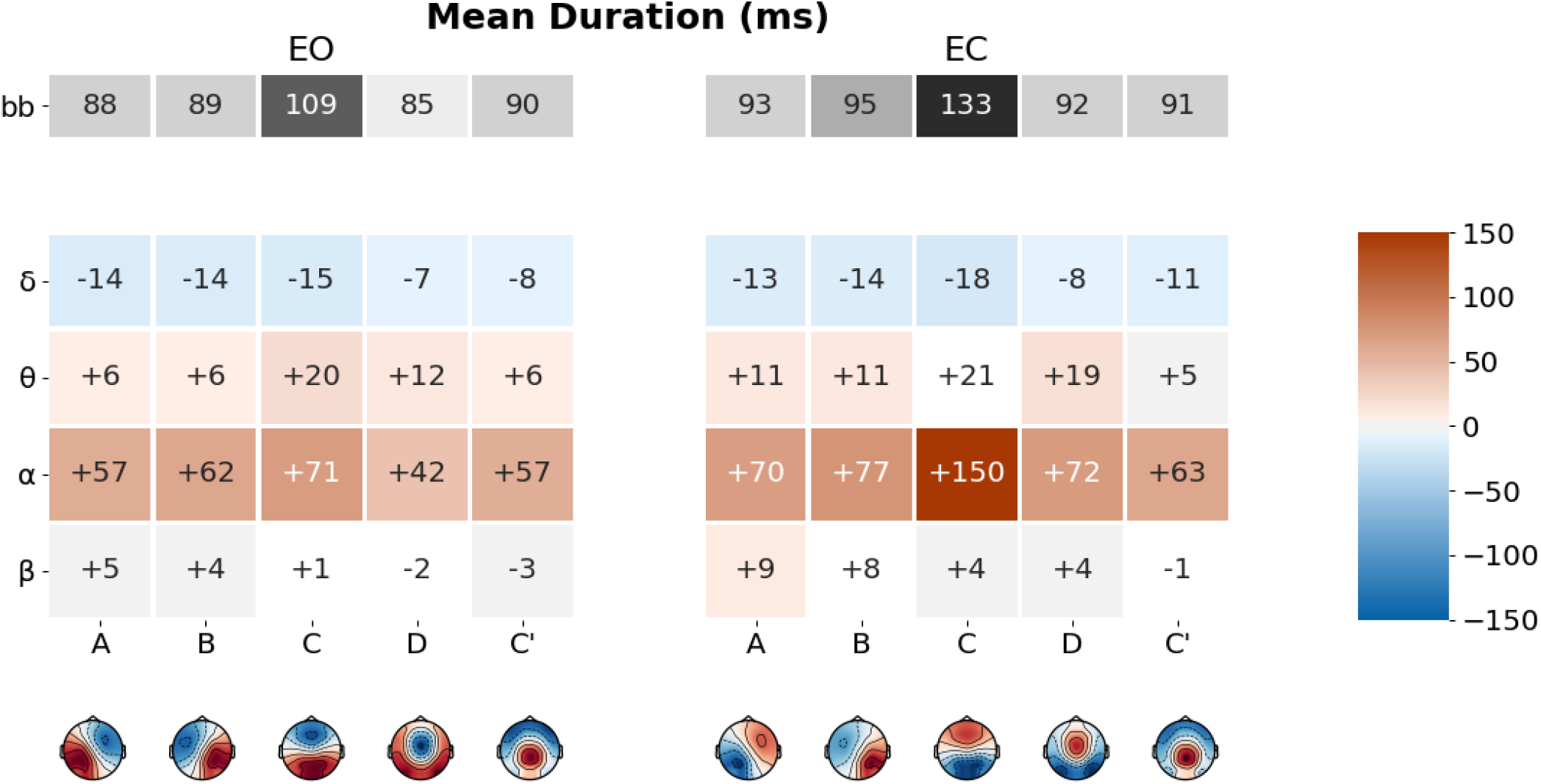
Microstate map differences in Mean Duration. (MeanDurs) between broadband and narrow-band filtered EEG, for eyes open (EO) and eyes closed (EC) conditions. The first row (in grey) represents the mean MeanDurs of the broadband segmentation. Coloured rows represent the mean MeanDurs difference between the narrowband (δ, θ, α, β) and broadband segmentations, for each microstate (A – C’). Significant differences have red/blue backgrounds, while non-significant ones have a white/grey background.

**Figure 7:**
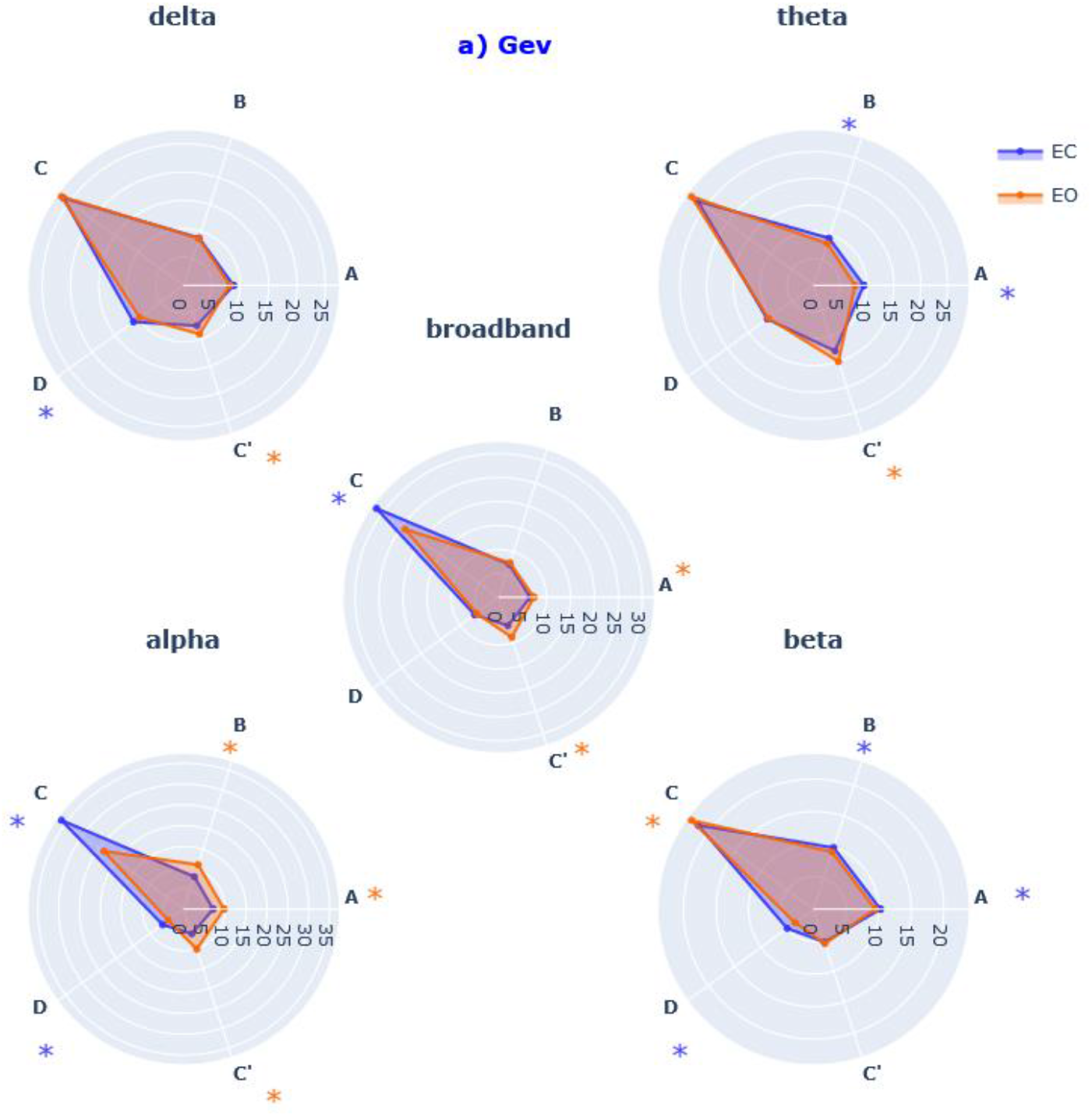
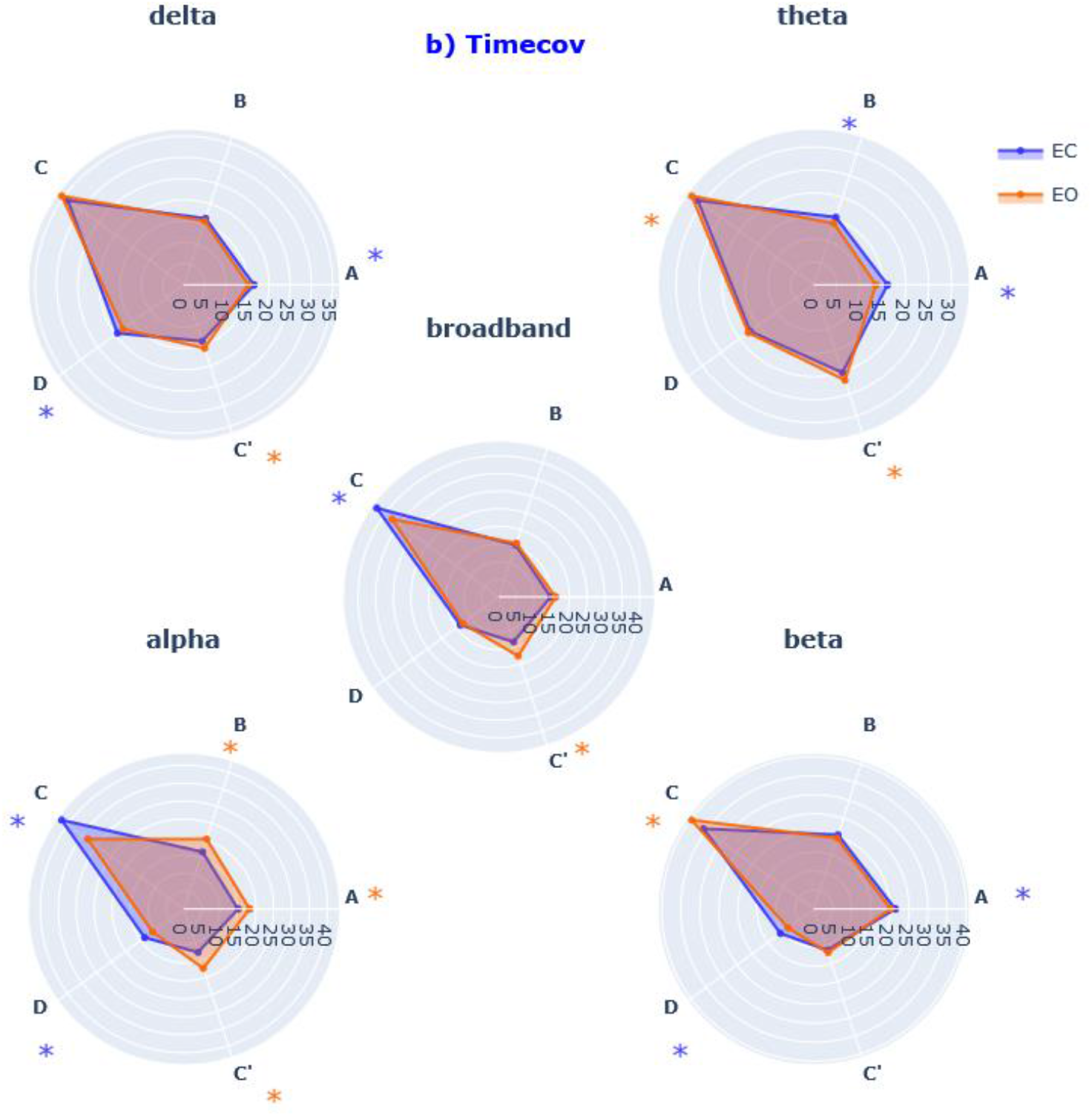
EO versus EC comparison using frequency specific microstate parameters. **a)** Mean global explained variance (Gev (%)),and **b)** mean time coverage (time coverage (%)) for each microstate (A – C’) within each frequency band (broadband, delta, theta, alpha, beta) for both eyes closed condition (EC, blue) and eyes opened condition (EO, red). Significance values are indicated from paired permutation test on mean between conditions: no star: 0.05 > p, *: p < 0.05, * color indicates the condition with highest value.

**Figure 8:**
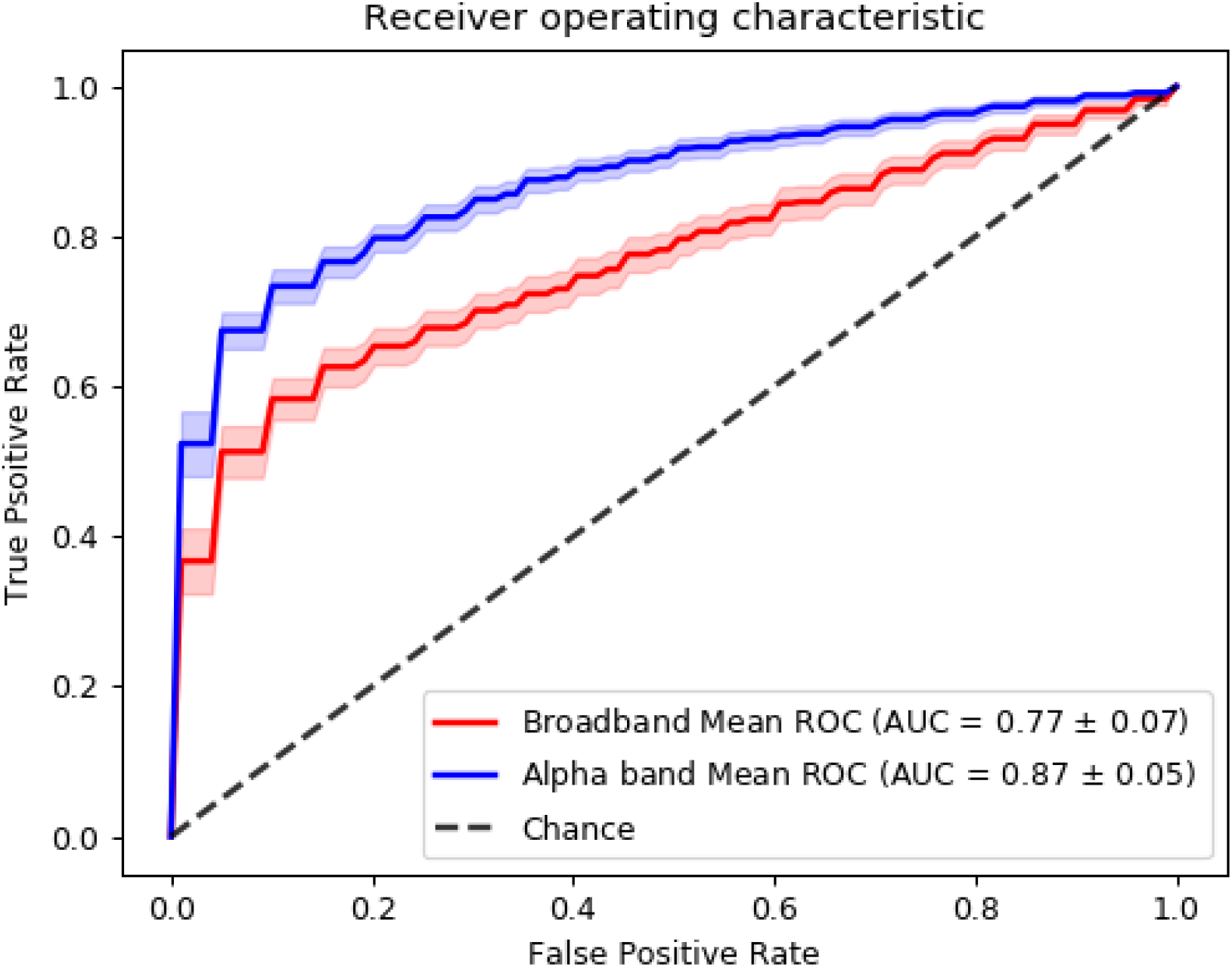
Classifying EO vs EC states using classical microstate measures. Binary classification performance, computed as the area under the curve (AUC) of the receiver-operating characteristic (ROC), for 2 support-vector machine models. Input features of the first model (red) were broadband MS parameters (see text) while the second model (blue) was evaluated on alpha-band MS parameters. Shaded areas represent 95% CI. Mean AUC ± standard deviation is reported in the legend.

Comparing the **delta** band segmentation to broadband, TimeCov was reduced for maps A (EO: - 3%, EC -0%), B (EO: - 2%, EC ns), and C (EO: 5%, EC: - 6 %), while it was increased for maps D (EO: + 5%, EC: + 4%) and C ‘(EO: +4, EC: +2%).

Small differences were generally observed between **theta** and broadband MS distributions, with the notable exception of a significant increase in map D (EO: + 5%, EC: + 7%) and decrease in map C (EO: - 2%, EC: - 4%), consistently found across behavioural states. For the **alpha** band, TimeCov was particularly greater compared to broadband for maps A (EO: + 2%, EC ns) and B (EO: + 4%, EC: + 1%)

Finally, comparing the **beta** band to broadband, TimeCov was increased for maps A (EO: +4%, EC: +5%, *p* < 0.05), B (EO: +4%, EC: +4%, *p* < 0.05), while it was reduced for maps D (EO: -4%, EC: -2%, *p* < 0.05) and C ‘(EO: -6%, EC: -6%, *p* < 0.05)

#### 3.3.2 Mean Duration (Mean Durs)

While analysing MS mean duration (MeanDurs) we first observed that MS were significantly shorter in **delta** compared to broadband across all maps: map A (EO: -14ms, EC: -13ms), map B (EO: -14ms, EC: -14ms), map C(EO:-15ms, EC: _-17ms, map D (EO: -7ms, EC: -8ms) and map C’ (EO: - 8ms, EC: -11ms).On the contrary **theta** segments were significantly longer than broadband ones for map A (EO: +6ms, EC: +11), map B (EO: +6ms, EC: +11ms) and map D(EO: -7ms, EC: -8ms). Similar but non-significant increases were also observed for maps C and C’.

**Notably**, all MS maps had significantly increased duration in **alpha** band compared to broadband: map A (EO: +57ms, EC: +70ms), map A (EO: +62ms, EC: +70ms) map C (EO: +71ms, EC: +150ms) map D (EO: +42ms, EC: +72ms) and map C’(EO: +57ms, EC: +63ms).

Less pronounced and less consistent differences were found for the **beta** band, which revealed a single significant increase of map A duration in the EC condition (EO: ns, EC: +9ms).

### 3.4 Within-frequency comparison of classical MS measures during eyes-open (EO) versus eyes-closed (EC)

Here, we directly compared EO vs EC conditions within each frequency band, and only relevant cases where narrow-band measures were salient compared to the broadband analysis are reported. Full results are documented in Table 3 and Suppl Figure 2 of the Supplementary Results.

#### 3.4.1 Global Explained Variance (Gev)

Across frequencies, map C’ explained more variance during EC than during EO for all bands (bb: d = 0.59, p<.05 | δ: d = 0.43, p<.05 | θ: d = 0.35, p<.05 | α: d = 0.35, p<.05 | β d = 0.67, ns) while map D expressed less variance(bb: d = -0.14, ns | δ: d = -0.07, ns | θ: d -0.29, p<.05 | α: d = - 0.42, p<.05 | β d = -0.73, p<.05). In contrast, the other MS maps had band-specific modulations: during EC map C explained relatively less variance in alpha (d = -1.0, p<.05) and broadband (d = -0.7, p<0.5),but expressed more variance in delta (d = 0.20, p<.O5) and beta (d = 0.21, p<.05). In addition, statistically significant effects were detected in the narrow-bands which were not evident in the broadband. For example, : the Gev of map B decreased from EC to EO (d = -0.31, p < 0.05) in in the beta band while no significant effect was found in broadband for this map.

#### 3.4.2 Time coverage (TimeCov)

Map C was relatively more prevalent in EC vs EO in alpha (d = -0.71, *p* < .05) and broadband (d = -0.48, *p* < .05), but the effect size was significantly stronger in the alpha band. In contrast, the opposite effect was observed in delta (d = 0.22, *p* < .05) and beta (d = 0.42, *p* < .05) bands, which showed increased coverage of map C during EO compared to EC. Diverging effects of time coverage in narrow-bands compared to broadband appeared across a number of MS maps and frequency bands, during the EO vs EC transition: *β*A –where TimeCov decreased prevalence (d = -0.20, *p* < .05), bbA and *β*C –TimeCov increased, respectively (d = 0.23, *p* < .05) and (d = 0.21, *p* <0 .05)., while bbC TimeCov decreased (d = -0.71, *p* < .05). Where non-significant differences between EO and EC conditions were found for θB coverage, *βB* (*d* =*-0*.*17, p*<.*05*) *and* θB (d = -0.35, p<.05) were significantly increased in the EC condition while αB (d = 0.57, p < .05) was decreased.

All in all, the EO vs EC-comparison revealed significant differences between the broadband and narrow-band segmentations, which were frequently map- and/or band-specific.

### 3.5 Statistical classification of EO vs EC behavioural states

As a proof-of-concept, we tested the applicability of band-specific microstates in the context of behavioural prediction, i.e. the binary discrimination of EO versus EC states using machine learning. Leveraging the well-known effect of alpha-band modulation during eye opening/closure (Barry et al. 2007) we hypothesised that alpha-band specific MS parameters would be stronger and more specific predictors of the EO/EC transition than those derived from the broadband segmentation. As we did not have any MS map specific hypotheses, we selected as features the basic set of classical MS measures from our study: 5 maps (A, B, C, D, C’) x 3 variables (Gev, TimeCov, MeanDurs). Then, using these 15 features, we applied a linear support vector machine (SVM) to classify EO vs EC recordings across all 203 subjects, separately for alpha-band and broadband models.

Following 10 fold cross-validation, we observed higher accuracy (i.e. sensitivity) for the SVM model tested with alpha-band MS parameters (80 ± 5%) than the one tested with broadband MS parameters (73 ± 6%). The superiority of the alpha-band model was reinforced by a separate analysis of the area under the curve (AUC) of the receiver-operating characteristic (ROC), which also incorporates the false-positive rate (i.e. specificity). Here alpha-band (AUC 87±5%) outperformed broadband (77±7%) by a full 10%. Finally, these differences were statistically significant for both accuracy (Cohen’s d=1.75, p<0 .001) and AUC (Cohen’s d=1.28, p <0 .001) demonstrating that a narrow-band MS segmentation may provide higher sensitivity/specificity in differentiating behavioural state relative to a broadband one.

## 4 Discussion

Historically, the first microstate (MS) analysis was applied by Lehmann and colleagues to narrowband (alpha) oscillations (Lehmann 1971), yet this “frequency-specific” approach appears to have been overlooked during the last decades of MS research in favour of decomposing broadband EEG signals (e.g. 2-40 Hz)(Michel and Koenig 2018; Pascual-Marqui et al. 1995). Hence, the present study specifically explored the MS characteristics of narrow-band EEG signals, their quantitative interrelationship, and whether they provide any novel information compared to course-grained broadband dynamics. This was done by first filtering the broadband EEG signal into several narrow-band frequencies (delta, theta, alpha, and beta), with the goal of comparing MS symbolic sequences and classical measures (explained variance, mean duration, time coverage) between them, as well as across different behavioural conditions (eyes open (EO) vs eyes closed (EC)).

### 4.1 The spatial dimension: microstate topographies

We first investigated whether analogous MS scalp topographies would be produced by segmenting broadband versus narrow-band EEG signals (including the alpha band (Milz et al. 2017)). Interestingly, we observed highly similar MS topographies (with minimum spatial correlations of *r* > 0.98) across all investigated broad- and narrow-band frequencies (broadband, delta to beta), as well as between EO/EC conditions. This is compatible with recent work by Brechet and colleagues (Bréchet et al. 2020), who observed that states of sleep and wake exhibited significantly different spectral content (e.g. delta vs beta power) but very similar MS maps. Moreover, these maps corresponded to the canonical (broadband) topographies previously described in the literature (Custo et al. 2017; Michel and Koenig 2018). It is therefore tempting to assume that identical neuronal sources are involved in generating the same topographies across frequencies. However, although different maps imply different generators (forward problem), same topographies do not necessarily imply identical generators (inverse problem). Due to the ill-posed nature of EEG signals (constructive and destructive electromagnetic fields), similar scalp potentials can still be generated by different underlying brain mechanisms (Helmholtz 1853). Hence, although we cannot unequivocally conclude that MS maps across the EEG spectrum are generated by the same brain sources operate, this would be the most probable and parsimonious interpretation. Moreover, we must juxtapose our findings with work from other groups (Musaeus et al. 2020) which applied a similar approach but didn’t necessarily find the same topographies across the EEG spectrum. From a methodological point of view, it should be kept in mind that narrow-band MS analysis does not *per se* require similar topographies between frequencies. In this case, although cross frequency comparisons would not be possible due to dissimilar maps, it would remain valid to study and quantify spatiotemporal MS parameters within each frequency band separately, for example, in the service of clinical biomarker discovery (Merrin et al. 1990). Reassuringly, the MS maps of our study replicate the ones derived from independent work utilising the same EEG dataset (Zanesco et al. 2020), further supporting the reproducibility of MS analysis despite methodological variations between studies (e.g. absence of resampling).

### 4.2 The temporal dimension: mutual information of microstate sequences

Milz and colleagues (Milz et al. 2017) recently proposed that alpha oscillations were the major component driving microstate dynamics. In general, adjusted mutual information (AMI) analyses reported in our work reveal low values (near or below 0.1) of information shared between the narrow-band segmentations, including alpha, and that of the broadband decomposition. However, consistent with the work of Milz and colleagues (Milz et al. 2017), the alpha-band during eyes-closed (EC) did indeed have the highest shared information with broadband (around 0.125). Importantly however, this relationship did not necessarily hold during eyes-open (delta being highest). This indicates specific narrow-band contribution(s) to broadband dynamics heavily depend on behavioural state. Moreover, if narrow-band(s) topographies were directly responsible for the origin of the spatial distribution of the broadband signal, one would expect much higher AMI values (at least 0.5) than those, we observed. In view of the results presented, it would be inaccurate to claim that alpha band, or any other narrow-band as the dominant source of broadband topographies.

In contrast, our results appear to support the ideas of Croce and colleagues (Croce et al. 2020), who suggested that broadband MS dynamics could not be extrapolated from one or a subset of EEG frequency bands. It remains unclear how the interaction of several narrow-band-components leads to a substantially different broadband MS decomposition. We speculate that this might stem from the fact that i) different narrow band signals could cancel each other at specific time points and ii) microstate assignment is non-linear given the winner-takes all approach.

Lastly and most intriguingly, no significant informational interrelations were found between the narrow-band topographical dynamics themselves (e.g. delta vs beta, theta vs alpha), indicating that each EEG band appears to have has its own independent dynamics. This may not be surprising, considering that spontaneous EEG oscillations have been reported to dynamically switch from a resting signatures (e.g. alpha) to task-specific active mode(s) dominated by theta (Ribary et al. 2017), beta (Fernández et al. 1995) or gamma activities (Hipp et al. 2011)). In this context, our observations of spatiotemporal independence between narrow-band EEG components support the operation of “oscillatory multiplexing” (Akam and Kullmann 2014) mechanisms in the cortex, whereby brain regions could combine different frequencies for integrating/segregating information across large-scale networks (Le Van Quyen 2011)

### 4.3 Classical microstate measures: explained variance, time coverage, and mean duration

#### Global Explained Variance (Gev)

Overall, MS segmentations of low-frequency bands (delta, theta) explained more of the global topographical variance compared to the classical broadband segmentaion. Since Gev is normalised by and therefore independent of global field power, this suggests that the goodness-of-fit (i.e Gev) of the five-map MS model is clearly greater in the delta/theta bands compared to broadband EEG. In higher frequencies, effects are more map-specific, as for example we observed an increase in Gev for the diagonal maps (A,B) and a decrease for midline (D,C’) ones..

#### Time Coverage (TimeCov)

Different patterns of MS temporal coverage were observed according to frequency. The delta and theta bands seem to exhibit greater prevalence of map D and decreased incidence of maps A, B, and C. This is remarkable insofar map D shares an intriguing overlap with the topography of the well-known frontal-midline theta-rhythm (fm-theta) (Scheeringa et al. 2008; Töllner et al. 2017). Conversely, beta band dynamics seem to favour the more frequent appearance of maps A and B in lieu of maps D and C’. The appearance of alpha-band MS maps, on the other hand, appear to be more state-dependent, as might be expected given the well-known expression of the posterior alpha rhythm during eyes-closed. Accordingly, there was a strong behavioural dissociation particularly for map C, which displayed a relatively greater temporal prevalence during EC compared to EO.

The presence of such “spectral fingerprints” suggests the existence of an affinity between distinct MS topographies and EEG bands suggests that different cortical generators (i.e. topographies) may be activated preferentially in certain frequencies (Groppe et al. 2013; Keitel and Gross 2016; Mellem et al. 2017).,.

#### Mean Duration (MeanDurs)

Microstates are defined as short periods of time during which the scalp electric field remains quasi-stable. Traditional microstate analysis does not suggest specific frequency filtering, thus resulting in various broadband filter settings across studies (Michel and Koenig 2018). Our findings demonstrate that temporally stable states (around 80ms or longer) are present within all classical EEG narrow-bands (i.e. delta to beta). It is established that such spatiotemporal structures do not appear for randomly shuffled EEG (Wackermann et al. 1993). For most EEG narrow-bands, mean MS durations were usually in the same range as the typically reported 70–120 ms, but often longer. For example, the average MS duration in the eyes open state was around 150 ms for the alpha-band, compared to 90 ms for broadband. It will be therefore interesting for future studies to examine the mechanistic links between the aggregated dynamics of the broadband microstates and those of frequency-specific modes.

### 4.4 Discriminating between behavioural states: eyes open versus eyes closed

Within each EEG narrow-band, between 8 (for theta) and 14 (for alpha) of the classical MS measures were found to be statistically significant. In comparison, classical MS measures derived from broadband EEG had a higher effect size for only one parameter (broadband microstate B mean duration). For all other 14 parameters, at least one narrow-band component showed a relatively stronger effect size.

Hence, the addition of the frequency dimension has the primary benefit of increasing both the number and the specificity of potential neural markers that could aid clinical prognosis or provide insight into brain mechanisms. We therefore conclude that the extra frequency dimension in itself leads to a more fine-grained decomposition of multichannel and multiplex EEG signals than the standard broadband analysis.

This is directly supported by our behavioural classification results, where we utilised alpha-band vs broadband MS parameters to predict EO vs EC states using the recordings of all 203 subjects. The significant boost in overall accuracy (alpha-band: 80% vs broadband: 73%) and area under the ROC curve nicely demonstrate that frequency-specific MS measures may provide higher behavioural predictive power than those derived from the broadband analysis. Although beyond the scope of this paper, we expect that this approach to provide advantages in other contexts, such as event-related potential (ERP) analyses or for discriminating between different clinical populations.

Interestingly, we found that in a few cases the narrow-band effects were opposite in directionality to the broadband results. Thus, limiting the analysis to only the latter could lead to incomplete (or even incorrect) interpretations of underlying brain dynamics.

## 5 Potential Limitations and Future Work

The current study may technically be considered exploratory, given the large number of tests that were carried out and in the absence of well-defined hypotheses, However, we carried out Bonferroni correction for all tests which may be considered the most conservative method for controlling for multiple comparisons. Several studies have thus far proposed explanations for the origins of broadband MS topographies (Britz et al. 2010). We feel it is still too early to make analogies or speculations between these results and those of the narrow-band dynamics. However, we believe that the application of the methodology proposed here may lead to valuable insights in order to more fully understand the underlying spectral tapestry of EEG microstates.

## 6 Conclusion

Ultimately, we report a number of important and novel findings between the classical broadband MS analysis, generally performed in the EEG field, and its application to more narrow frequency bands relevant to cortical oscillatory activities. In a nutshell, it appears that each canonical EEG frequency band possesses its own independent spatiotemporal dynamics, whilst the relative prevalence of MS topographies themselves differ across frequencies. Moreover, it appears that broadband dynamics cannot be appropriately explained by the simple summation of individual narrow-band frequency components.

Analysis of narrow-band MS parameters revealed spatial and temporal characteristics that both converged and diverged from broadband MS findings. At the very least, our results indicate that narrow-band analysis is justified as complementary to the usual broadband MS analysis. A narrow-band decomposition into frequencies more specific for cortical oscillatory activity could not only advance and consolidate findings in clinical disorders e.g. (Merrin et al. 1990) (Musaeus et al. 2020), but also enable a better understanding of the organization and functioning of large-scale brain systems..

## Data and Code Availability Statement

The data and code supporting our results are openly available in the OSF repository and can be found at: https://osf.io/bfsge/ (DOI 10.17605/OSF.IO/BFSGE).

Other 3rd party software and datasets used in this study can be found by following citations in the paper.

## Data availability statement

The data and code supporting our results are openly available in the OSF repository and can be found at: https://osf.io/bfsge/

## Acknowledgments

The study was supported by the Swiss National Science Foundation (NCCR Synapsy grant No. 51NF40 – 185897 and grant No. 320030_184677) to CMM.

We also would like to thank the Mind-Body-Emotion group at the Max Planck Institute for Human Cognitive and Brain Sciences for all the work they have done to make their dataset public.

## Conflict of Interest Statement

The authors declare that the research was conducted in the absence of any commercial or financial relationships that could be construed as a potential conflict of interest.

## Notes

https://osf.io/bfsge/

### Competing Interest Statement

The authors have declared no competing interest.

### Summary of Updates

General revision of the article and figures, added section "Statistical classification of EO vs EC behavioural states".

## References

Akam, Thomas, and Dimitri M. Kullmann. 2014. ‘Oscillatory Multiplexing of Population Codes for Selective Communication in the Mammalian Brain’. Nature Reviews Neuroscience 15(2):111–22. doi: 10.1038/nrn3668.

Armstrong, Richard A. 2014. ‘When to Use the B Onferroni Correction’. Ophthalmic and Physiological Optics 34(5):502–8.

Babayan, Anahit, Miray Erbey, Deniz Kumral, Janis D. Reinelt, Andrea MF Reiter, Josefin Röbbig, H. Lina Schaare, Marie Uhlig, Alfred Anwander, Pierre-Louis Bazin, and others. 2019. ‘A Mind-Brain-Body Dataset of MRI, EEG, Cognition, Emotion, and Peripheral Physiology in Young and Old Adults’. Scientific Data 6:180308.

Barry, Robert J., Adam R. Clarke, Stuart J. Johnstone, Christopher A. Magee, and Jacqueline A. Rushby. 2007. ‘EEG Differences between Eyes-Closed and Eyes-Open Resting Conditions’. Clinical Neurophysiology 118(12):2765–73. doi: 10.1016/j.clinph.2007.07.028.

Bouckaert, Remco R. n.d. ‘Choosing between Two Learning Algorithms Based on Calibrated Tests’. 8.

Bréchet, Lucie, Denis Brunet, Gwénaël Birot, Rolf Gruetter, Christoph M. Michel, and João Jorge. 2019. ‘Capturing the Spatiotemporal Dynamics of Self-Generated, Task-Initiated Thoughts with EEG and FMRI’. Neuroimage 194:82–92.

Bréchet, Lucie, Denis Brunet, Lampros Perogamvros, Giulio Tononi, and Christoph M. Michel. 2020. ‘EEG Microstates of Dreams’. Scientific Reports 10(1):17069. doi: 10.1038/s41598-020-74075-z.

Britz, Juliane, Dimitri Van De Ville, and Christoph M. Michel. 2010. ‘BOLD Correlates of EEG Topography Reveal Rapid Resting-State Network Dynamics’. NeuroImage 52(4):1162–70. doi: 10.1016/j.neuroimage.2010.02.052.

Brunet, Denis, Micah M. Murray, and Christoph M. Michel. 2011. ‘Spatiotemporal Analysis of Multichannel EEG: CARTOOL’. Computational Intelligence and Neuroscience 2011.

Croce, Pierpaolo, Angelica Quercia, Sergio Costa, and Filippo Zappasodi. 2020. ‘EEG Microstates Associated with Intra-and Inter-Subject Alpha Variability’. Scientific Reports 10(1):1–11.

Custo, Anna, Dimitri Van De Ville, William M. Wells, Miralena I. Tomescu, Denis Brunet, and Christoph M. Michel. 2017. ‘Electroencephalographic Resting-State Networks: Source Localization of Microstates’. Brain Connectivity 7(10):671–82.

D’Croz-Baron, David F., Mary Baker, Christoph M. Michel, and Tanja Karp. 2019. ‘EEG Microstates Analysis in Young Adults With Autism Spectrum Disorder During Resting-State’. Frontiers in Human Neuroscience 13:173. doi: 10.3389/fnhum.2019.00173.

Fernández, Thalía, Thalía Harmony, Mario Rodríguez, Jorge Bernal, Juan Silva, Alfonso Reyes, and Erzsébet Marosi. 1995. ‘EEG Activation Patterns during the Performance of Tasks Involving Different Components of Mental Calculation’. Electroencephalography and Clinical Neurophysiology 94(3):175–82. doi: 10.1016/0013-4694(94)00262-J.

Gramfort, Alexandre, Martin Luessi, Eric Larson, Denis A. Engemann, Daniel Strohmeier, Christian Brodbeck, Roman Goj, Mainak Jas, Teon Brooks, Lauri Parkkonen, and others. 2013. ‘MEG and EEG Data Analysis with MNE-Python’. Frontiers in Neuroscience 7:267.

Groppe, David M., Stephan Bickel, Corey J. Keller, Sanjay K. Jain, Sean T. Hwang, Cynthia Harden, and Ashesh D. Mehta. 2013. ‘Dominant Frequencies of Resting Human Brain Activity as Measured by the Electrocorticogram’. NeuroImage 79:223–33. doi: 10.1016/j.neuroimage.2013.04.044.

Helmholtz, H. von. 1853. ‘Ueber Einige Gesetze Der Vertheilung Elektrischer Strome in Korperlichen Leitern, Mit Anwendung Auf Die Thierisch-Elektrischen Versuche (Schluss.)’. Annalen Der Physik 165(7):353–77.

Hipp, Joerg F., Andreas K. Engel, and Markus Siegel. 2011. ‘Oscillatory Synchronization in Large-Scale Cortical Networks Predicts Perception’. Neuron 69(2):387–96. doi: 10.1016/j.neuron.2010.12.027.

Javed, Ehtasham, Pierpaolo Croce, Filippo Zappasodi, and Cosimo Del Gratta. 2019. ‘Hilbert Spectral Analysis of EEG Data Reveals Spectral Dynamics Associated with Microstates’. Journal of Neuroscience Methods 325:108317.

Keitel, Anne, and Joachim Gross. 2016. ‘Individual Human Brain Areas Can Be Identified from Their Characteristic Spectral Activation Fingerprints’. PLoS Biology 14(6):e1002498.

Koenig, Thomas, and Daniel Brandeis. 2016. ‘Inappropriate Assumptions about EEG State Changes and Their Impact on the Quantification of EEG State Dynamics’. Neuroimage 125:1104–6.

Le Van Quyen, Michel. 2011. ‘The Brainweb of Cross-Scale Interactions’. New Ideas in Psychology 29(2):57–63. doi: 10.1016/j.newideapsych.2010.11.001.

Lehmann, D. 1971. ‘Multichannel Topography of Human Alpha EEG Fields’. Electroencephalography and Clinical Neurophysiology 31(5):439–49.

Mellem, Monika S., Sophie Wohltjen, Stephen J. Gotts, Avniel Singh Ghuman, and Alex Martin. 2017. ‘Intrinsic Frequency Biases and Profiles across Human Cortex’. Journal of Neurophysiology 118(5):2853–64.

Merrin, Edward L., Patricia Meek, Thomas C. Floyd, and Enoch Callaway III. 1990. ‘Topographic Segmentation of Waking EEG in Medication-Free Schizophrenic Patients’. International Journal of Psychophysiology 9(3):231–36.

Michel, Christoph M., and Thomas Koenig. 2018. ‘EEG Microstates as a Tool for Studying the Temporal Dynamics of Whole-Brain Neuronal Networks: A Review’. NeuroImage 180:577–93. doi: 10.1016/j.neuroimage.2017.11.062.

Milz, Patricia, Roberto D. Pascual-Marqui, Peter Achermann, Kieko Kochi, and Pascal L. Faber. 2017. ‘The EEG Microstate Topography Is Predominantly Determined by Intracortical Sources in the Alpha Band’. Neuroimage 162:353–61.

Musaeus, Christian S., Knut Engedal, Peter Høgh, Vesna Jelic, Arjun R. Khanna, Troels Wesenberg Kjær, Morten Mørup, Mala Naik, Anne-Rita Oeksengaard, Emiliano Santarnecchi, and others. 2020. ‘Changes in the Left Temporal Microstate Are a Sign of Cognitive Decline in Patients with Alzheimer’s Disease’. Brain and Behavior e01630.

Pascual-Marqui, Roberto D., Christoph M. Michel, and Dietrich Lehmann. 1995. ‘Segmentation of Brain Electrical Activity into Microstates: Model Estimation and Validation’. IEEE Transactions on Biomedical Engineering 42(7):658–65.

Pedregosa, F., G. Varoquaux, A. Gramfort, V. Michel, B. Thirion, O. Grisel, M. Blondel, P. Prettenhofer, R. Weiss, V. Dubourg, J. Vanderplas, A. Passos, D. Cournapeau, M. Brucher, M. Perrot, and E. Duchesnay. 2011. ‘Scikit-Learn: Machine Learning in Python’. Journal of Machine Learning Research 12:2825–30.

Ribary, Urs, S. M. Doesburg, and L. M. Ward. 2017. ‘Unified Principles of Thalamo-Cortical Processing: The Neural Switch’. Biomedical Engineering Letters 7(3):229–35. doi: 10.1007/s13534-017-0033-4.

Ros, Tomas, Bernard J. Baars, Ruth A. Lanius, and Patrik Vuilleumier. 2014. ‘Tuning Pathological Brain Oscillations with Neurofeedback: A Systems Neuroscience Framework’. Frontiers in Human Neuroscience 8. doi: 10.3389/fnhum.2014.01008.

Scheeringa, René, Marcel C. M. Bastiaansen, Karl Magnus Petersson, Robert Oostenveld, David G. Norris, and Peter Hagoort. 2008. ‘Frontal Theta EEG Activity Correlates Negatively with the Default Mode Network in Resting State’. International Journal of Psychophysiology 67(3):242–51. doi: 10.1016/j.ijpsycho.2007.05.017.

Schulman, Joshua J., Robert Cancro, Sandlin Lowe, Feng Lu, Kerry D. Walton, and Rodolfo R. Llinás. 2011. ‘Imaging of Thalamocortical Dysrhythmia in Neuropsychiatry’. Frontiers in Human Neuroscience 5. doi: 10.3389/fnhum.2011.00069.

Töllner, Thomas, Yijun Wang, Scott Makeig, Hermann J. Müller, Tzyy-Ping Jung, and Klaus Gramann. 2017. ‘Two Independent Frontal Midline Theta Oscillations during Conflict Detection and Adaptation in a Simon-Type Manual Reaching Task’. The Journal of Neuroscience 37(9):2504–15. doi: 10.1523/JNEUROSCI.1752-16.2017.

Van De Ville, D., J. Britz, and C. M. Michel. 2010. ‘EEG Microstate Sequences in Healthy Humans at Rest Reveal Scale-Free Dynamics’. Proceedings of the National Academy of Sciences 107(42):18179–84. doi: 10.1073/pnas.1007841107.

Vinh, Nguyen Xuan, Julien Epps, and James Bailey. 2010. ‘Information Theoretic Measures for Clusterings Comparison: Variants, Properties, Normalization and Correction for Chance’. The Journal of Machine Learning Research 11:2837–54.

Wackermann, J., D. Lehmann, CM Michel, and WK Strik. 1993. ‘Adaptive Segmentation of Spontaneous EEG Map Series into Spatially Defined Microstates’. International Journal of Psychophysiology 14(3):269–83.

Zanesco, Anthony P., Brandon G. King, Alea C. Skwara, and Clifford D. Saron. 2020. ‘Within and Between-Person Correlates of the Temporal Dynamics of Resting EEG Microstates’. NeuroImage 211:116631. doi: 10.1016/j.neuroimage.2020.116631.

