## Supplementary figures and images for "Beyond broadband: towards a spectral decomposition of EEG microstates"

### Suppl Figure 1

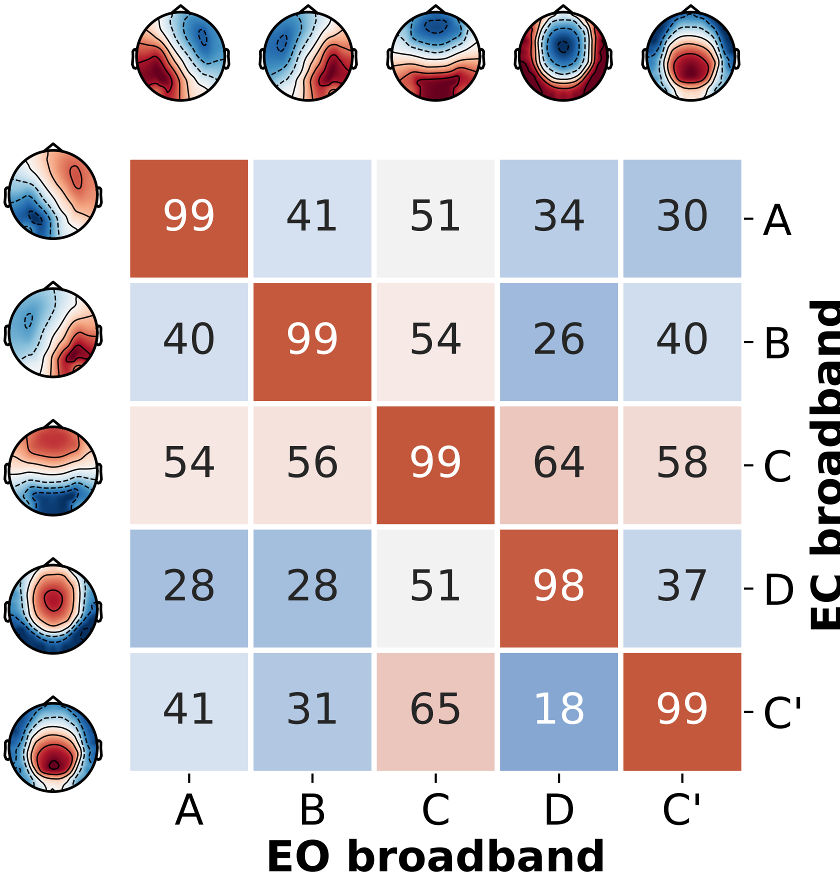

### Suppl Figure 2

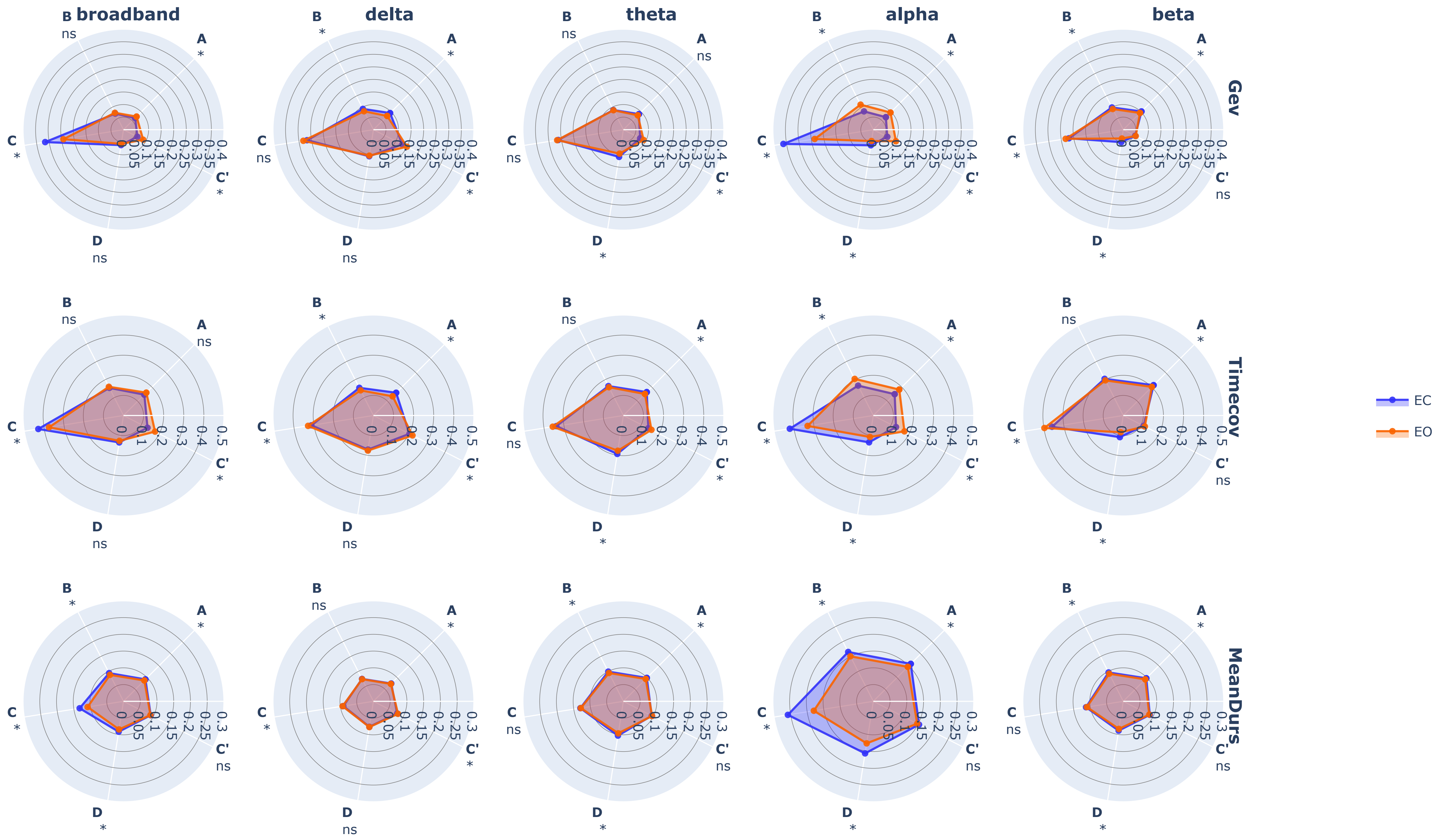
